# Optical Torque Calculations and Measurements for DNA Torsional Studies

**DOI:** 10.1101/2024.05.29.596477

**Authors:** Yifeng Hong, Fan Ye, Jin Qian, Xiang Gao, James T. Inman, Michelle D. Wang

**Affiliations:** Department of Electrical and Computer Engineering, Cornell University, Ithaca, NY 14853, USA; Howard Hughes Medical Institute, Cornell University, Ithaca, NY 14853, USA; Department of Physics & LASSP, Cornell University, Ithaca, NY 14853, USA

## Abstract

The angular optical trap (AOT) is a powerful instrument for measuring the torsional and rotational properties of a biological molecule. Thus far, AOT studies of DNA torsional mechanics have been carried out using a high numerical aperture oil-immersion objective, which permits strong trapping, but inevitably introduces spherical aberrations due to the glass-aqueous interface. However, the impact of these aberrations on torque measurements is not fully understood experimentally, partly due to a lack of theoretical guidance. Here, we present a numerical platform based on the finite element method to calculate forces and torques on a trapped quartz cylinder. We have also developed a new experimental method to accurately determine the shift in the trapping position due to the spherical aberrations by using a DNA molecule as a distance ruler. We found that the calculated and measured focal shift ratios are in good agreement. We further determined how the angular trap stiffness depends on the trap height and the cylinder displacement from the trap center and found full agreement between predictions and measurements. As further verification of the methodology, we showed that DNA torsional properties, which are intrinsic to DNA, could be determined robustly under different trap heights and cylinder displacements. Thus, this work has laid both a theoretical and experimental framework that can be readily extended to investigate the trapping forces and torques exerted on particles with arbitrary shapes and optical properties.

**SIGNIFICANCE:** We developed a simulation platform based on the finite element method for force and torque calculation for particles in an angular optical trap (AOT), with considerations of tightly focused Gaussian beam, spherical aberrations, and optically anisotropic particles. Experimental measurements of focal shift ratio, force, and torque under multiple conditions were in good agreement with predictions from the simulations. We also demonstrated that intrinsic DNA torsional properties can be robustly measured under different AOT measurement conditions, strongly validating our simulations and calibrations. Our platform can facilitate trapping particle design for single-molecule assays using the AOT.

## INTRODUCTION

Optical tweezers (also known as optical traps) utilize light to manipulate and measure the dynamics of individual biological molecules and are powerful tools for studying protein-DNA interactions (1,2). Due to the inherent helical structure of DNA, rotational motions are integral to fundamental processes that take place over DNA. As DNA-based motor proteins translocate, they rotate DNA (3,4). While optical tweezers have been highly successful in measuring biological forces (5-8), they cannot measure torques and rotational motions. The angular optical trap (AOT) overcame this limitation, enabling simultaneous measurements of forces and torques of a biological molecule (9-12). The AOT has been used to measure the torque generated by RNA polymerase (13,14), the torque required to destabilize a nucleosome (15), the torsional modulus of a single and braided chromatin fiber (16), and the force-torque relation of the DNA phase diagram (11,17-24).

Unlike conventional optical tweezers that trap a dielectric microsphere, the AOT traps a nanofabricated birefringent cylinder with the bottom of the cylinder functionalized for the attachment of a DNA molecule (Fig. 1) (11). The cylinder is typically made of quartz with its extraordinary axis perpendicular to the cylindrical axis and is optically trapped with its cylindrical axis aligned to the direction of the light propagation. The cylinder is rotated via rotation of the incoming light’s linear polarization, and the optical torque exerted on the cylinder is measured via the change in the angular momentum of the light after the light interacts with the cylinder.

**FIGURE 1.**
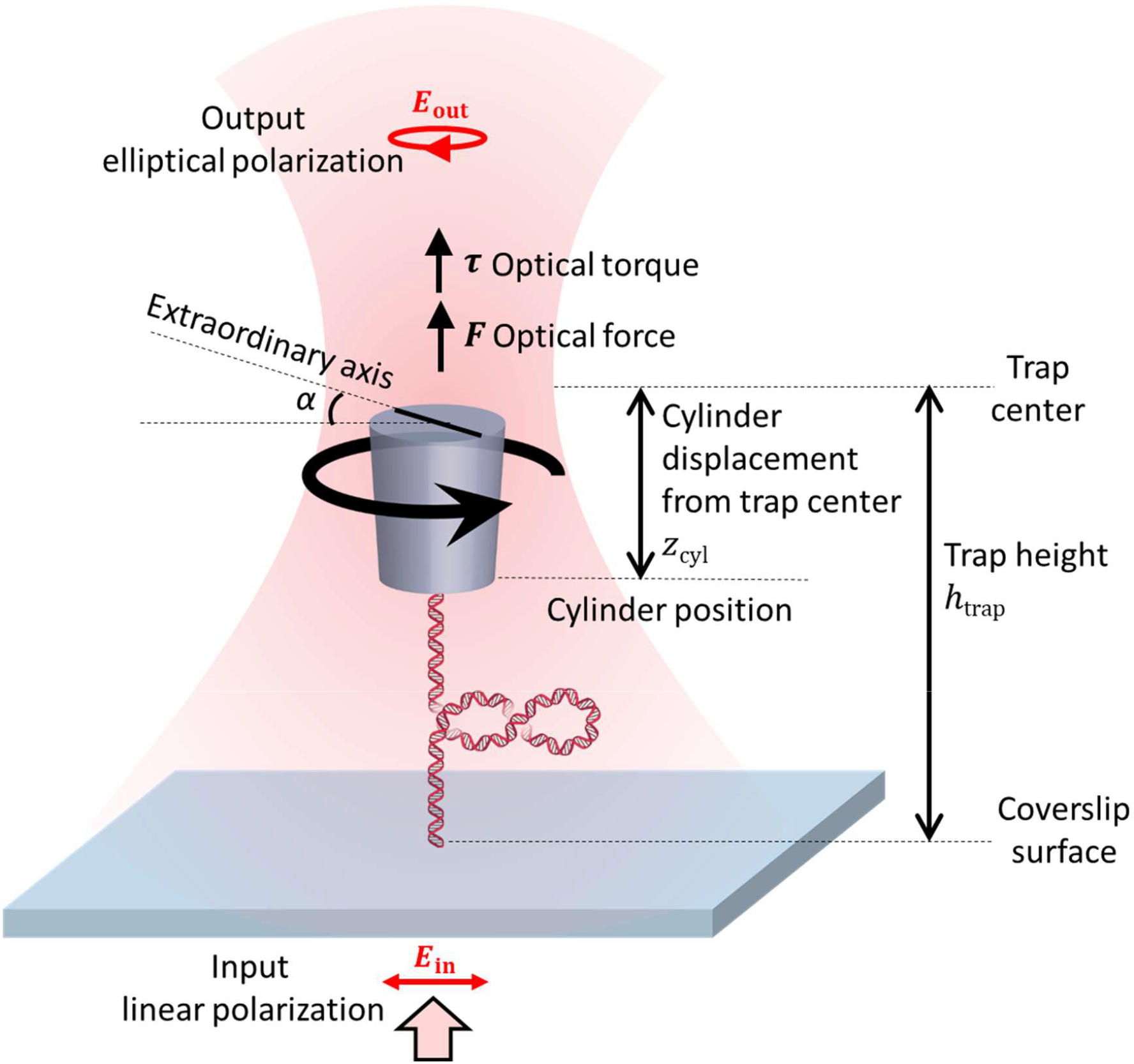
Experimental configuration on the AOT. The trapping laser is initially linearly polarized and becomes slightly elliptically polarized after exerting a torque on a nanofabricated quartz cylinder, which is tethered to a DNA molecule and held in the trap. The trap height *h*_trap_ (the distance of the trap center from the surface) is controlled by the surface height via a piezo stage. If the cylinder is displaced (*z*_cyl_) from the trap center by the DNA, then the trap exerts an axial force on the cylinder to resist this displacement. If the cylinder is used to add turns to the DNA, the trap exerts a torque on the cylinder, resulting in a slight misalignment angle α between the cylinder’s extraordinary axis and the trapping beam polarization. Both the optical force and optical torque are directly measured.

An AOT requires a high numerical aperture (NA) objective to focus the trapping laser beam to a tight spot near the specimen plane. Thus far, DNA torque measurements and behaviors of DNA-based motor proteins have primarily relied on an oil-immersion objective (11,13-24). While an oil-immersion objective permits efficient trapping, it also inevitably introduces spherical aberrations at the trapping region due to the refractive index mismatch when the trapping beam passes through the interface between the glass coverslip and the aqueous solution of a sample chamber (Fig. 2a). The resulting spherical aberrations lead to an axial shift of the beam waist (focal shift) with a concurrent distortion of the trapping beam.

**FIGURE 2.**
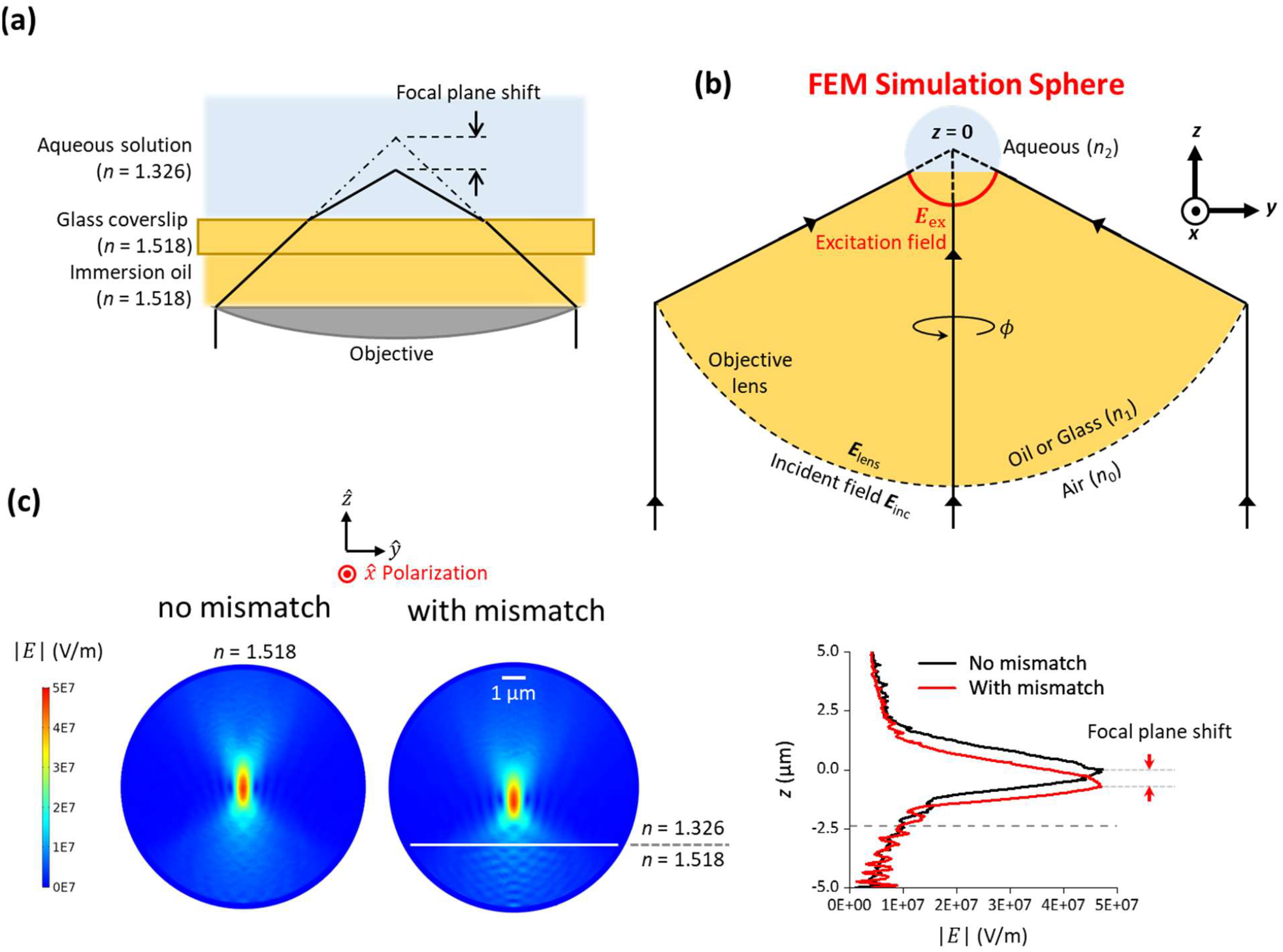
Simulation of a tightly focused Gaussian beam near a glass-aqueous interface. (a) A cartoon illustrating the spherical aberrations induced by the glass-aqueous interface based on ray optics. The dotted dash lines intercept the expected focal plane in absence of the index mismatch, while the solid lines intercept the shifted focal plane. (b) A sketch illustrating the numerical approach of calculating the tightly focused Gaussian beam for the FEM simulations. (c) Simulated E-field distributions of the focused beam without/with the index mismatch at 1 W laser power at the specimen plane.

These changes could potentially compromise trapping beam quality and complicate force and torque calibrations. Previous studies have examined the axial shift of the equilibrium trapping position of a trapped particle due to the glass-aqueous interface (25-31). However, theoretical guidance on the angular stiffness dependence on the focal depth has thus far been lacking.

In this work, we have developed a numerical method to explicitly consider how a laser beam is focused through the glass-aqueous interface by a high NA oil-immersion objective.

Using this method, we have calculated the trap center shift, the force, and the torque on a trapped quartz cylinder using the finite element method (FEM) via COMSOL Multiphysics. We then developed a method to accurately measure axial trap center shift based on DNA stretching and experimentally determined the axial linear trap stiffness and angular trap stiffness as a function of the trap height. We found a good agreement between simulations and measurements. Our theoretical understanding and experimental validation have enabled robust measurements of DNA torsional properties, despite the spherical aberrations. This work lays down a numerical framework that is suitable for a broad range of applications that examine force and torque properties of trapping particles with arbitrary shape, size, and anisotropy, when trapped by a distorted Gaussian beam.

## MATERIALS AND METHODS

### Fabrication of functionalized quartz cylinders

The quartz cylinders were fabricated by adapting protocols we previously established (11,19,23). We started the fabrication process from a 4-inch X-cut quartz wafer (Precision Micro-Optics). The top surface of the quartz wafer was initially functionalized with 3-aminopropyltriethoxysilane (Sigma, 440141) solution (11). The functionalized quartz cylinders were ultimately yielded from the top-down fabrication based on deep ultraviolet (DUV) lithography and reactive ion etching (RIE) (19). Finally, the functionalized area of the quartz cylinders was covalently attached with streptavidin (Agilent, SA10-10) via glutaraldehyde (Sigma, G5882) linkage to anchor DNA for single-molecule measurements. Cylinders used in this work have the following dimensions as measured by scanning electron microscope (mean ± SD): 473 ± 35 nm bottom diameter, 589 ± 43 nm top diameter, and 1016 ± 45 nm height (*N* = 24).

### DNA template preparation

#### Torsionally unconstrained DNA templates

The 6,546 bp and 11,516 bp single-labeled DNA were prepared via PCR from plasmid pMDW133 (sequence available upon request). The 6,546 bp DNA was made with the forward 5’-digoxigenin-labeled primer 5’-GCCACCTGACGTCTAAGAAACCATTATTATCA and the reverse 5’-biotin-labeled primer 5’-GCTGGTCCTTTCCGGCAATCAGG. The 11,516 bp DNA was made with the forward 5’-digoxigenin-labeled primer 5’-GCCACCTGACGTCTAAGAAACCATTATTATCA and the reverse 5’-biotin-labeled primer 5’-ACGAGACGAAAAAACGGACCGCGTTTG. The final products were purified via gel extraction.

#### Torsionally constrained DNA template

The 6,481 bp DNA multi-labeled DNA was prepared via PCR from plasmid pMDW133 as described previously (20).

### Data acquisition and analysis

All the AOT experiments were performed at room temperature (23 °C). Each sample chamber used for measurements was assembled from a nitrocellulose-coated coverslip and a Windex-cleaned glass slide (16). The force calibration of cylinders was performed in phosphate buffered saline (PBS, Invitrogen AM9625: 137 mM NaCl, 2.7 mM KCl, 8 mM Na_2_HPO_4_, 2 mM KH_2_PO_4_, pH 7.4). All the other experiments were performed in PBS with 0.6 mg/mL β-casein (Sigma, C6905). DNA-tethered cylinders for the AOT measurements were prepared by following a previously reported assay (19).

#### DNA stretching

The DNA stretching experiment was carried out in two steps. In the first step, the DNA was stretched with no feedback at a constant laser power until *z*_cyl_ reached 300 nm. Then, the DNA continued to be stretched under a constant *z*_cyl_ mode (feedback by laser power) until the stretching force reached 20 pN (6). Data were recorded at a sampling rate of 400 Hz.

#### Linear trap stiffness dependence on trap height h_trap_

The axial linear trap stiffness and the axial position detector sensitivity at different trap heights were carried out based on a previously developed DNA unzipping method (26) at laser powers of 50.4 mW and 67.2 mW at the specimen plane. The unzipping speed was 50 nm/s. Data were recorded at 400 Hz.

### Angular stiffness dependence on h_trap_

The measurement of torque at different trap heights was carried out with un-tethered cylinders based on previous protocols (12,30,31). At a given trap height, we first calibrated the angular sensitivity. Then, the calibrated angular sensitivity was used to convert the angular power spectral density data for angular stiffness determination at each trap height (32). The sampling rates of these measurements were 10 kHz.

### Angular stiffness dependence on axial cylinder displacement z_cyl_

The measurement of torque at different *z*_cyl_ was carried out with torsionally unconstrained DNA-tethered cylinders under different forces (0.4 pN, 0.75 pN, 1.5 pN, 2.2 pN, 3.0 pN, 3.7 pN) at an estimated 8.4 mW laser power at the specimen plane, where the beam polarization was rapidly rotated at 500 turn/s. The sampling rate for this fast winding experiment was 10 kHz.

### DNA twisting

The DNA twisting experiments were carried out under a constant-force mode of 3 pN (19) (feedback by piezo) at laser powers of 10.5 mW and 42 mW at the specimen plane, respectively. The quartz cylinder was rotated at 2 turn/s to add turns to the torsionally constrained DNA attached underneath. Data were recorded at a sampling rate of 400 Hz. The torque data were smoothed with sliding windows of 2 s.

## RESULTS AND DISCUSSION

### Simulations of a highly focused beam through a glass-aqueous interface

We have developed a method to determine how a high NA objective focuses a Gaussian beam through a glass-aqueous interface via a combination of analytical and simulation methods. As illustrated in Fig. 2b, we started by analytically calculating the electric field of a far-field, linearly polarized paraxial Gaussian beam (*λ* _0_ = 1064 nm) focused and apertured by an aplanatic optical lens in the absence of any glass-aqueous interface near the laser focus (33) (Supplementary Information). Here, this lens is an oil-immersion objective: Nikon NA = 1.3 with an aperture filling factor 0.98 in our design (30,31), which is index-matched with the glass coverslip. We obtained the electric field at a spherical surface immediately after passing through the objective, which focuses all the rays towards the center of a sphere (*x, y, z*) = (0, 0, 0). We then propagated all rays analytically towards the focus to a reference sphere until the sphere reached a radius of 5 μm. The electric field at the surface of this sphere then served as the excitation field for the spherical finite-element method (FEM) simulation near the focal plane by COMSOL. This treatment minimized the COMSOL computational resource. Inside this spherical simulation space, we introduced the glass-aqueous interface and determined how the presence of this interface altered the field distribution near the laser focus (Fig. 2c; Fig. S1). The distorted laser beam was focused slightly below the center of the sphere. We placed a quartz cylinder in the aqueous solution near the laser focus and computed the optical force – 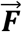 and optical torque 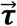 on the cylinder:

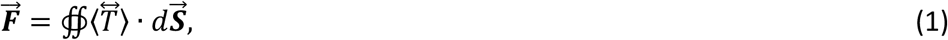

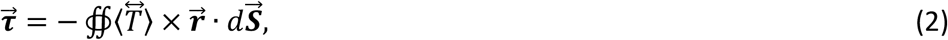

where 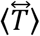 is the time-averaged Maxwell stress tensor and 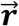 is the vector pointing from the cylinder’s geometric center to its surface. For an optically anisotropic trapping particle (for quartz, *n*_e_ = 1.5428 and *n*_o_ = 1.5341 at 1064 nm (34)), the Maxwell stress tensor can be written as (35)

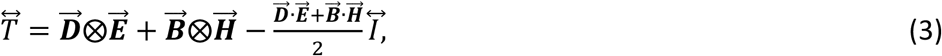

where 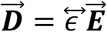 is the electric displacement, 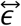 is the permittivity tensor of the material (Supplementary Information), 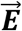 is the electric field, 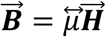 is the magnetic field, 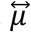 is the material permeability tensor, 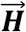 is the magnetic intensity, ⨂ is the dyadic product, and 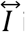 is the unit tensor.

Previously, an index-mismatched interface was considered in simulations using the transition matrix method (36), which is computationally efficient but challenging to be extended to inhomogeneous and anisotropic particles (37), whereas the FEM method is more flexible (38). Our approach captures all the critical features induced by spherical aberrations on an optically anisotropic particle as discussed below.

### Focal shift ratio determination

The presence of the glass-aqueous interface shifts the laser focus towards the interface (*h*_focus_) (Fig. 2a), resulting in a corresponding axial shift of the equilibrium trapping position of an untethered cylinder (*h*_trap_), which is also referred to as the trap center (Fig. 1; Fig. 3a). This is characterized by the focal shift ratio *f*_s_, defined as the ratio of the axial trap height change Δ*h*_trap_ over the axial surface change Δ*z*_surf_ (2,26,32):

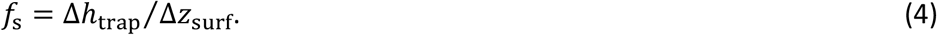

In the absence of spherical aberrations, *f*_s_ = 1 so that when the surface is moved up by Δ*z*_surf_, there is a corresponding decrease in the trap height by the exact same amount. When spherical aberrations are present, *f*_s_ < 1.

**FIGURE 3.**
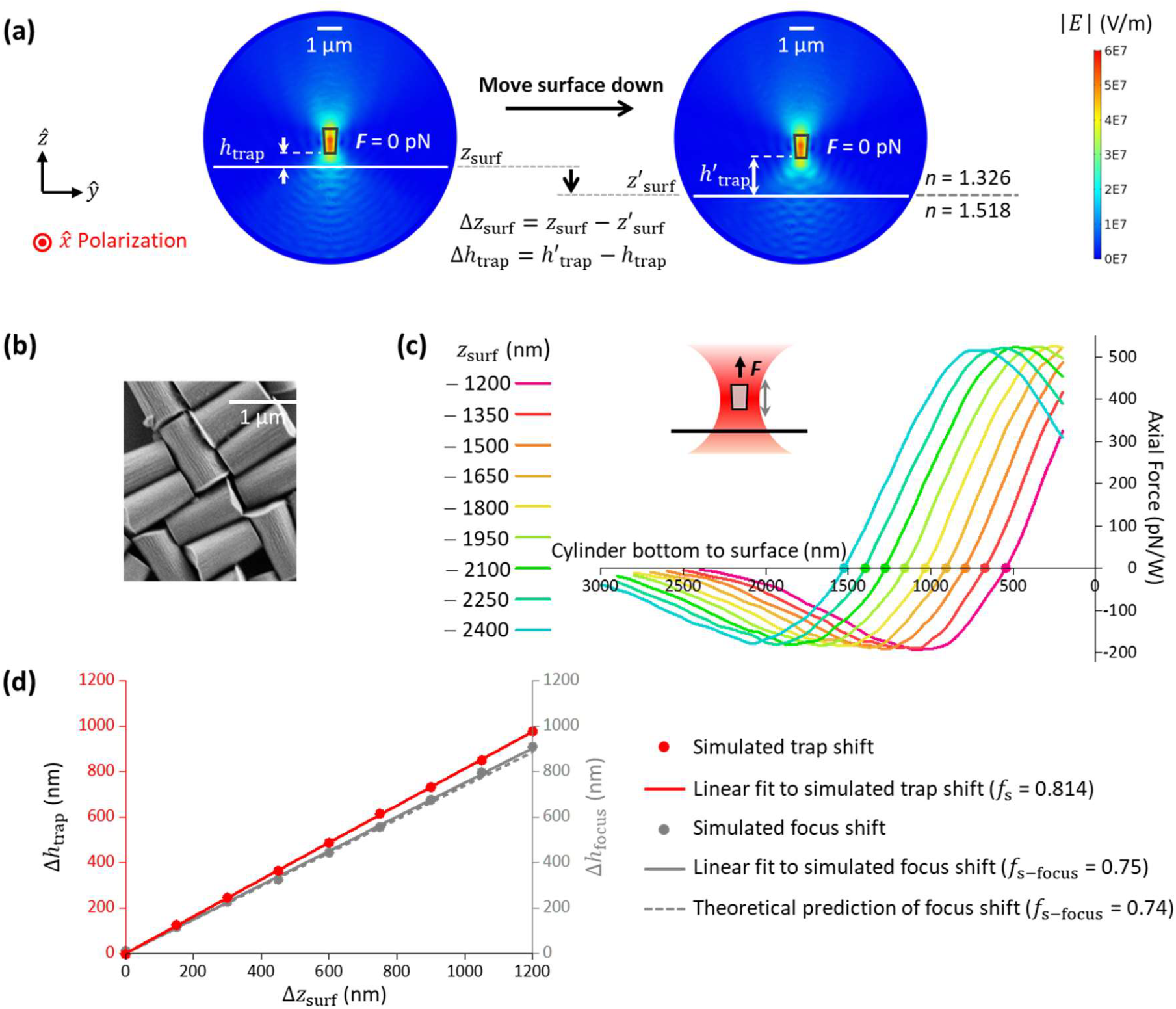
Simulations of the trapping force and focal shift ratio for a trapped quartz cylinder. The simulations were carried out with the cylinder’s extraordinary axis aligned with the input trapping laser polarization so there is no net torque on the cylinder. (a) E-field distributions of a trapped quartz cylinder at its equilibrium position at two surface positions. *z*_surf_ indicates the coverslip surface position. *h*_trap_ is defined as the distance between the cylinder’s bottom surface to the coverslip surface. (b) An SEM image of nanofabricated quartz cylinders. (c) Trap height. Simulated axial force profile of a quartz cylinder along z-axis at different *z*_surf_ positions. The equilibrium position of the cylinder occurs when the net force is zero (indicated as a dot on each curve), which indicates the trap height *h*_trap_. When *z*_surf_ = 0, the surface is located at the center of the simulation sphere which is also the focus of the laser beam. The power indicated is at the specimen plane. (d) Focal shift. A linear fit to Δ*h*_trap_ versus Δ*z*_surf_ relation gives the focal shift ratio: *f*_s_ = 0.814 (solid line). For comparison, we also show how the laser focus changes with surface displacement based on our simulations and a theoretical prediction.

To calculate this ratio, we performed simulations on a cylinder with the same dimensions as those we have fabricated (Fig. 3b) so that the simulation results can be directly compared with measurements. At a given surface height, we calculated the trapping force versus cylinder position relative to the surface (Fig. 3c). The trap center is located where the cylinder experiences no net force. We repeated this calculation at different surface heights to obtain the relation of Δ*h*_trap_ versus Δ*z*_surf_ (Fig. 3d). A linear fitting to this relation yielded *f*_s_ = 0.814 ± 0.003 (fit ± fit error). This value is somewhat smaller than the value of *n*_water_/*n*_glass_ = 0.87 expected from a simple paraxial ray approximation of the laser focus (39), suggesting that such a simple approximation does not fully capture the axial beam distortion caused by spherical aberrations, but is larger than the focal shift ratio of the laser focus obtained from our simulations: 0.75 ± 0.01 (fit ± fit error) (Fig. 3d). In addition, the focal shift of the laser focus from our simulations is in agreement with a prediction from a previous theoretical study (0.74) (40), providing some confirmation for the accuracy of our simulations. These simulation results indicate that the cylinder is pushed further downstream of the laser focus as the trap height increases.

To compare the simulations with measurements, we first performed simulations to determine the force on a cylinder that is attached to the surface as the surface is moved through the beam focus (Fig. 4a). We also conducted corresponding force measurements using the same configuration. We found that the measured force is in good agreement with the force from simulations if we assume the laser transmission coefficient of our objective is 0.42. This value is comparable to those previously reported for an oil immersion objective (41). For all subsequent comparisons, we have kept this value unchanged.

**FIGURE 4.**
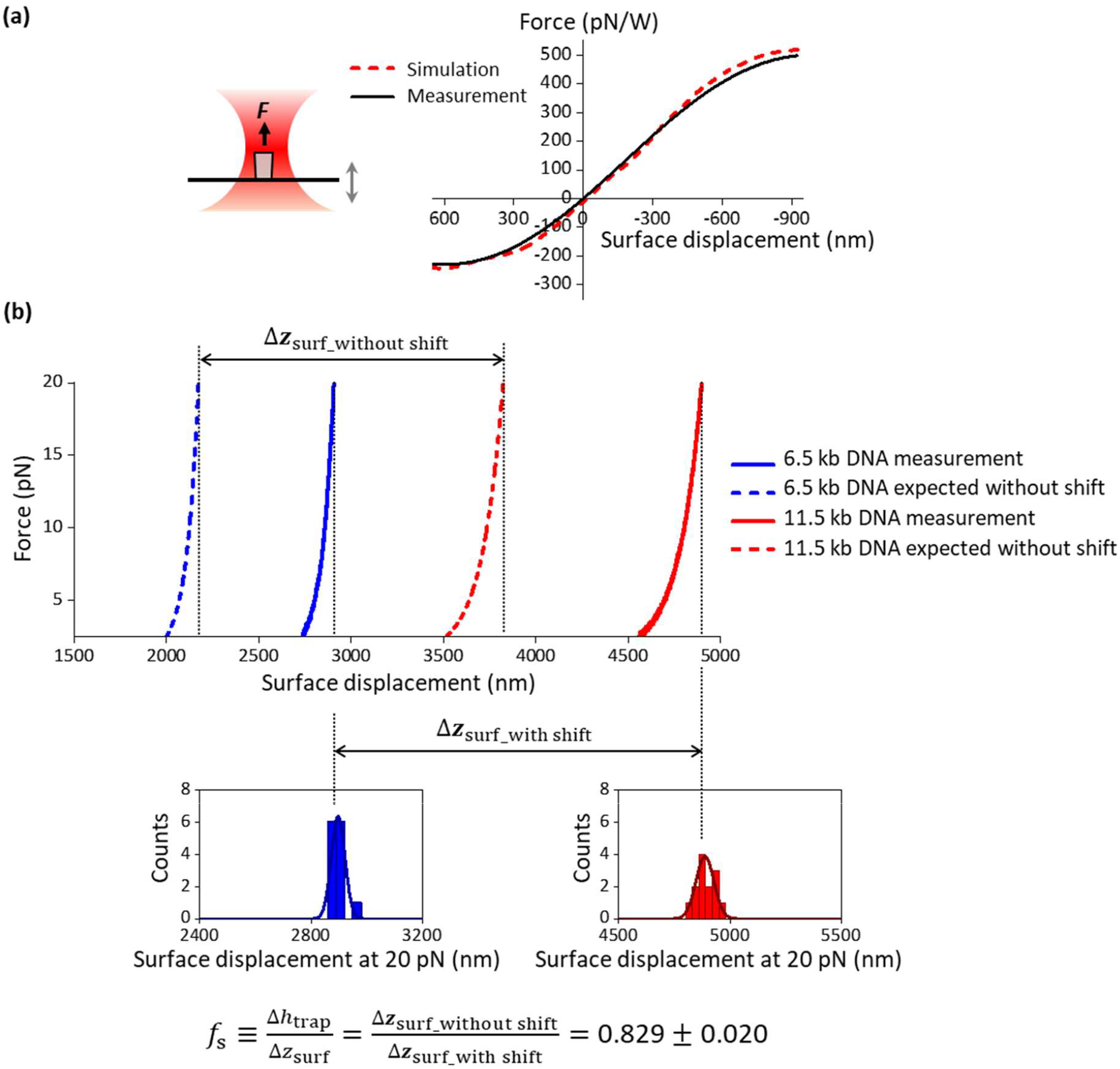
Experimental validation of force and focal shift ratio simulations. (a) Measured force on a surface-attached quartz cylinder versus the coverslip surface movement and its comparison with simulation. To make the force simulations feasible in COMSOL, the bottom surface of the cylinder was placed 10 nm above the surface. The measured force profile is the averaged result of *N* = 13 cylinders. The experimental laser power in the specimen plane was estimated by assuming that the objective has a transmission coefficient of 0.42. (b) Experimental determination of the focal shift ratio. The top panel shows example traces of stretching dsDNA molecules of two different lengths to obtain the force versus coverslip surface displacement relation. Δ***z***_surf_without shift_ is the expected difference in the coverslip surface positons at 20 pN for DNA molecules of two different lengths if there is no focal shift. Δ***z***_surf_with shift_ is the measured difference in the coverslip surface positons in the presence of the focal shift. The focal shift ratio of the trap is thus obtained from the ratio of these two differences. The SD in the *f*_s_ is determined via error propagation from measured extensions of the molecules of the two different lengths at 20 pN.

To compare the calculated focal shift ratio with the measured value, we need a method to accurately determine this ratio. Previously, two methods have been used to estimate this ratio. One method is based on the Fabry-Pérot effect that originates from the beam interference between the bottom of the cylinder and the coverslip surface (25,32). This approach can provide an estimate for the focal shift ratio, but requires the plane-wave assumption which needs to be validated for a high NA objective (42). An alternative approach utilizes DNA unzipping force signatures (26,32), but this approach requires prior knowledge of ssDNA elasticity parameters which are sensitive to DNA sequence and ionic conditions (43).

Here, we present an alternative experimental method that is more robust. This method uses dsDNA as a distance ruler and relies on the fact that when a dsDNA molecule is stretched to ∼ 20 pN, its extension is close to its contour length of 0.338 nm/bp and is insensitive to variations in dsDNA elasticity parameters under various buffer conditions (6). Consequently, the dsDNA extension at this force is an excellent distance ruler for focal shift ratio determination.

In Fig. 4b (top plot), we measured the force as a function of the coverslip surface displacement for two different lengths of dsDNA molecules (6.5 kb and 11.5 kb). For each molecule, to reach a given force value, the coverslip surface had to be displaced by a much larger distance than the expected extension from the force-extension relation of the modified Marko-Siggia worm-like-chain model (6). This discrepancy informs the focal shift ratio.

Although this method can in principle be implemented using dsDNA of a single length, using dsDNA molecules of two different lengths allows more accurate focal shift determination. This method critically depends on accurate determination of the coverslip position when the DNA reaches zero extension, but this determination can have a small but non-negligible offset on the order of 20 nm (31). In addition, there might be small extension offsets arising from extension contributions of the end anchors of the DNA molecules. To minimize the contribution of these offsets to the focal shift determination, we instead examine the difference in the coverslip positions at 20 pN of the two different molecule lengths to obtain the focal shift ratio *f*_s_ (Fig. 4b). Using this difference method, we determined *f*_s_ = 0.829 ± 0.020 (mean ± SD). Using this focal shift ratio, we found the resulting DNA force-extension curve shows an excellent agreement with the expectation (Fig. S2). As a comparison, we found that the focal shift ratio experimentally obtained based on the Fabry-Pérot effect yielded *f*_s_ = 0.829 ± 0.003 (mean ± SD) (Fig. S3). Thus, focal shift ratios obtained using the DNA stretching method and the Fabry-Pérot method are in excellent agreement.

It is worth noting that because force and displacement calibrations require prior knowledge of the focal shift ratio, an initial estimate of this ratio must be used for these calibrations, which then allow the determination of a more accurate focal shift ratio. Thus the final focal shift ratio must be obtained iteratively.

### Angular stiffness dependence on trap height and cylinder displacement

The trapping beam becomes more distorted as the beam is further focused into the aqueous solution, and this distortion weakens the axial linear trap stiffness at an increased trap height (26). Indeed, we found that the measured linear trap stiffness decreases by about 3.4% per μm of trap height (Fig. S4). However, it is unclear as to whether this distortion could weaken the angular trapping stiffness at an increased trap height. To investigate this possibility, for a given trap height, we simulated the torque versus α, the angle of deviation of the cylinder’s extraordinary axis from the linear polarization of the input beam (Fig. 1; Fig. 5a). Experimentally, we calibrated the angular stiffness at different trap heights. The simulated and measured relations are in excellent agreement. Surprisingly, the angular trap stiffness shows nearly no dependence on the trap height over a few microns of the trap height as demonstrated by both the simulations and measurements. This lack of dependence suggests that angular properties are rather insensitive to the beam distortions, simplifying the angular stiffness calibrations.

**FIGURE 5.**
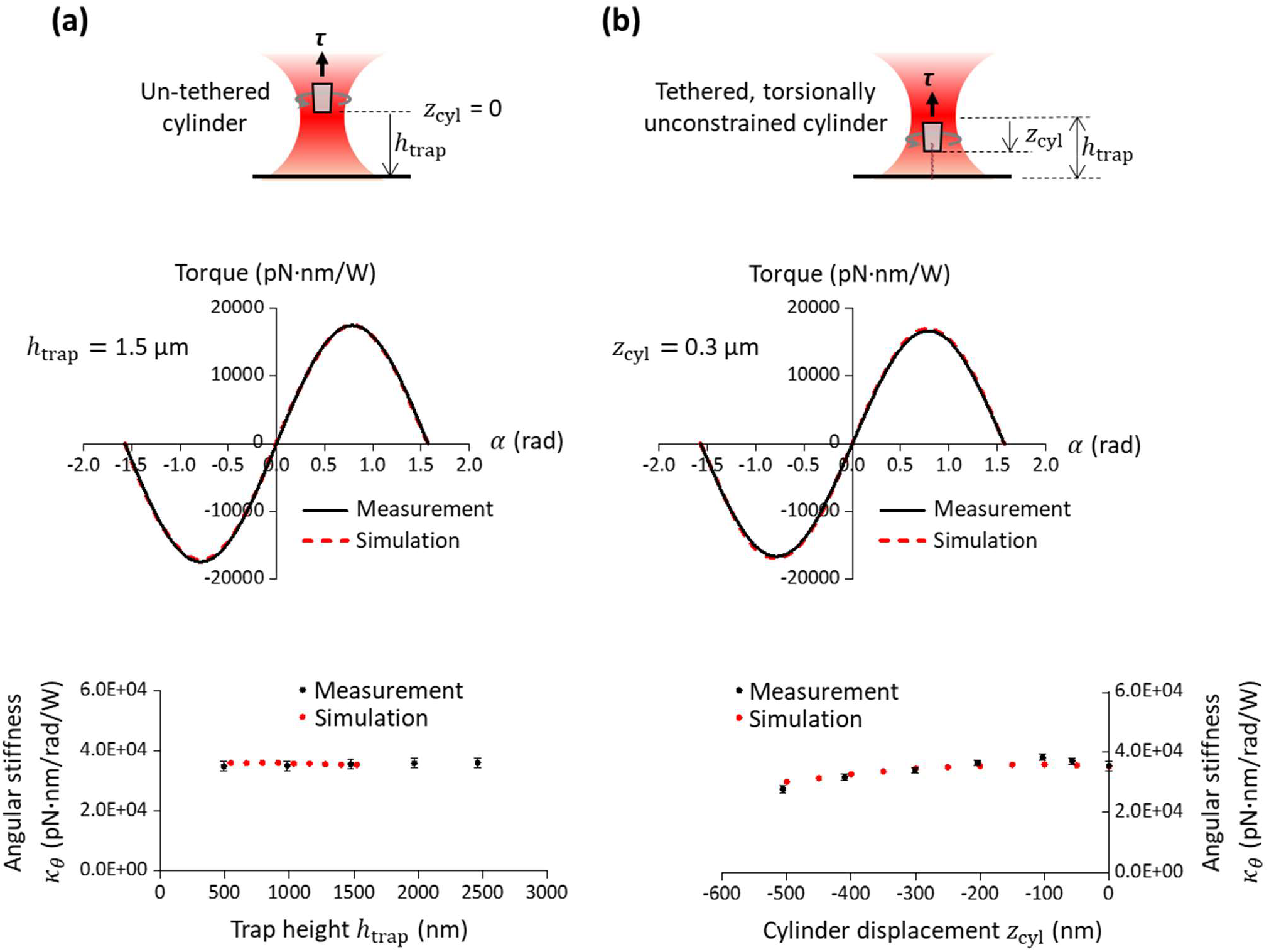
Experimental validation of the torque simulations. (a) Angular stiffness dependence on trap height *h*_trap_. The top panel shows a schematic experimental configuration for angular stiffness measurements using an un-tethered cylinder. The middle panel shows the measured torque versus angular displacement α at *h*_trap_ = 1.5 µm (averaged from *N* = 11 cylinders), a comparison with simulation. This relation yields the angular stiffness. The bottom panel shows the measured angular stiffness dependence on the trap height with error bars being SEMs, and a comparison with simulation. (b) Angular stiffness dependence on cylinder displacement *z*_cyl_. The top panel shows a schematic experimental configuration for angular stiffness measurements using a DNA tethered cylinder. The DNA was tethered to the cylinder without torsional constraint so that the cylinder was displaced from the trap but was allowed to freely rotate. The middle panel shows the measured torque versus angular displacement α at *z*_cyl_ = 0.3 µm (averaged from *N* = 10 cylinders) and a comparison with simulation. The bottom panel shows the measured angular stiffness dependence on cylinder displacement with error bars being SEMs, and a comparison with simulation.

In addition, during experimental DNA torsional measurement (Fig. 1), the AOT is often used to apply a force on the DNA as the DNA pulls the cylinder downward from the trap center. During this process, the cylinder will move through a beam profile with an axially varying intensity, which may lead to possible changes in the angular stiffness. This consideration exists even in the absence of a glass-aqueous interface. To investigate this possibility, we simulated the torque versus α relation at different *z*_cyl_ values (Fig. 5b) at a fixed trap height. The simulation results show a very slight increase in the angular stiffness for *z*_cyl_ from 0 to -100 nm, followed by a gradual decrease in the angular stiffness with further cylinder displacement, consistent with the cylinder’s overlap with the beam profile as it moves across the beam focus and then moves away from it.

To evaluate this dependence experimentally, we pulled the cylinder from the trap center by a desired *z*_cyl_ using a DNA molecule that was tethered to the cylinder bottom but without torsional constraint. We then measured the torque versus α relation by rapidly rotating the polarization, where the cylinder was quasi-stationary during the measurement (10,12). The resulting measured angular stiffness is almost an exact match to that from the simulation and also shows a similar trend with an increase in the downward cylinder displacement.

### DNA torsional measurements

The ultimate verification for simulations and calibrations should be their performance during experimental applications of DNA torsional measurements. The goal is to robustly determine DNA torsional properties under different measurement conditions. Here, we measured DNA torsional response under a constant force (Fig. 6), which has clearly defined extension and torque characteristics (11,16,19,21,23,44,45): when turns are added to DNA, the DNA extension does not have significant change while torque in DNA increases linearly until the DNA buckles to extrude a plectoneme; after DNA buckling, DNA extension decreases while the torque remains constant. Since these features are intrinsic to DNA, they should be independent of the trap height and the cylinder displacement used in the measurements.

**FIGURE 6.**
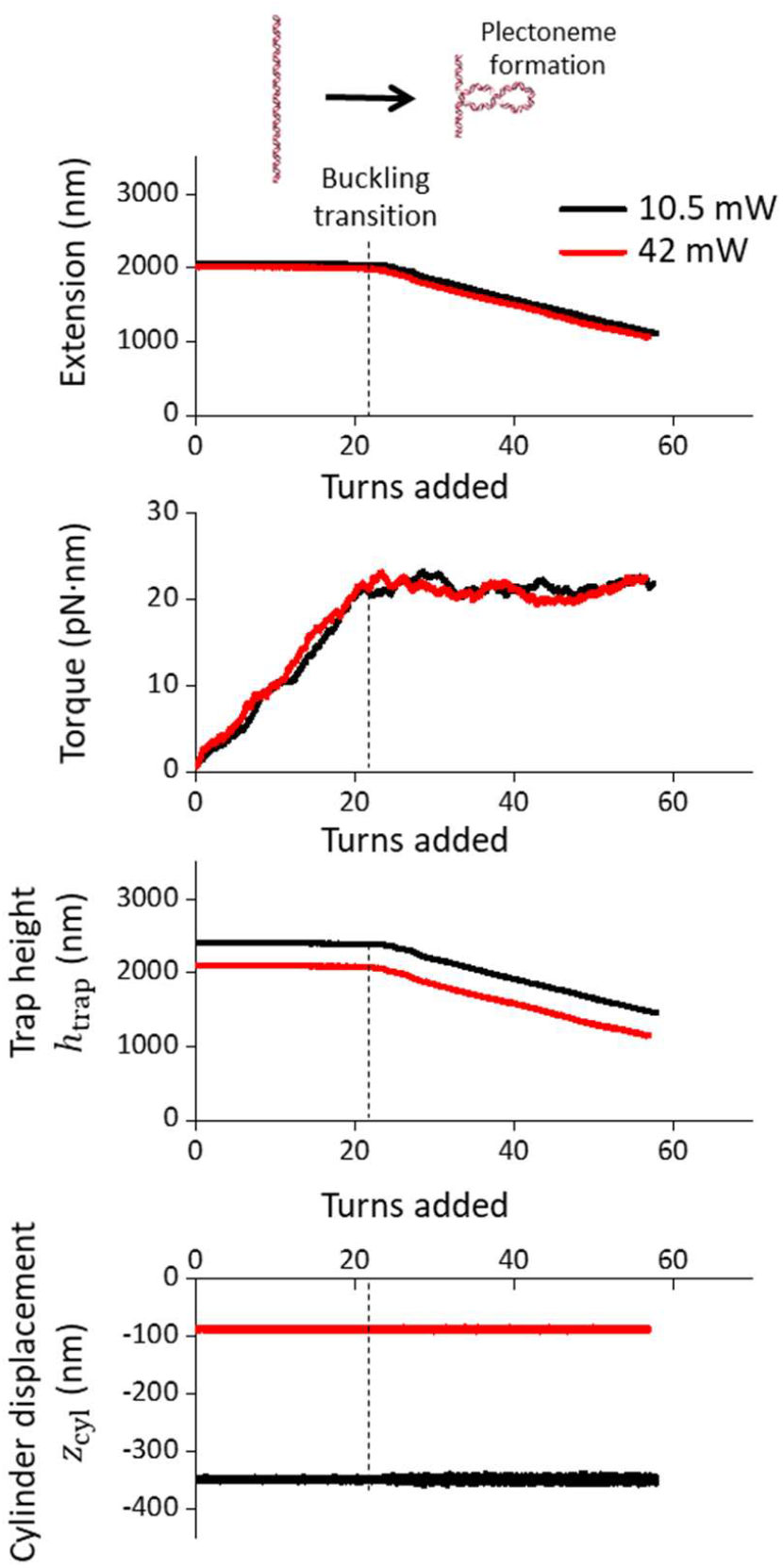
DNA torsional measurements. A 6.5 kb DNA molecule was torsionally constrained between the cylinder and the coverslip surface (Fig. 1). Using the AOT, we held the DNA under a constant force of 3 pN and added turns to DNA. For a given laser power, the force clamp was achieved by keeping the cylinder displacement *z*_cyl_ constant via the modulation of the trap height using a piezo stage. Consequently, the lower the laser power, the larger the cylinder displacement. Traces shown are the average of multiple DNA tethers (*N* = 7 for 10.5 mW laser power at the specimen plane; *N* = 7 for 42 mW at the specimen plane).

As shown in Fig. 6, we introduced sufficient turns to buckle the DNA where the extension decreased significantly, necessitating a corresponding decrease in the trap height. As expected, the measured buckling torque remained constant when the trap height decreased after DNA buckling. To test the performance of torque measurements under different *z*_cyl_ values, we carried out the experiments under two different laser powers that led to *z*_cyl_ = 357 nm and 89 nm. Again, the torque signals remained the same for both *z*_cyl_ values. Thus, these data provide strong support for the validity of our simulations and experimental methods.

## CONCLUSION

We have developed a simulation platform via COMSOL that addresses all core features of the AOT based on a high NA oil-immersion objective by considering a tightly focused Gaussian beam, spherical aberrations, nanofabricated anisotropic cylinders, and force/torque generation. We also developed the method of dsDNA stretching for accurate focal shift ratio determination. Our simulations and experimental validations unravel the dependences of the AOT torque measurements on trap height and cylinder displacement, which were further supported by consistent DNA torsional measurements. Although the use of the oil-immersion objective introduces spherical aberrations, we show that the primary complication is the need to accurately determine the focal shift ratio. Interestingly, the angular stiffness of the trap is rather insensitive to these aberrations when experiments are performed within a few microns of the glass-aqueous interface.

Our finding of the focal shift determined by using the DNA stretching method being consistent with that from the Fabry-Pérot method suggests that the plane-wave assumption for the Fabry-Pérot method is reasonable even for a high NA objective. This confirmation validates the use of the Fabry-Pérot method, which does not require DNA substrates, is more compatible with a broader range of trapping configurations, and is more straightforward in measurements and data analysis.

Our work facilitates understanding of the AOT instrumentation considerations and single-molecule data interpretation. Although this work focuses on quartz cylinders, it can be readily extended as a general model for optical force and torque calculation regardless of the size, shape, composition, and orientation of a trapped object (46-48). It has valuable potential for broader usage in developing and exploring novel nanotechnologies coupled with the AOT.

## DATA AND CODE AVAILABILITY

We have provided the source Mathematica code via a Github repository (https://github.com/WangLabCornell/2024_Hong_et_al). This Mathematica code numerically calculates the electric field of a Gaussian beam after passing through a high NA objective and outputs the electric field at a spherical boundary and generates inputs for the commercial software COMSOL.

## AUTHOR CONTRIBUTIONS

Y. H. and F. Y. performed simulations. Y. H. fabricated the quartz cylinders. Y. H. performed cylinder calibrations and measurements with J.Q.’s help. Y. H. analyzed data. X. G. and J. T. I. optimized the angular optical trap setup. M. D. W. supervised the project.

## ACKNOWLEDGMENTS

We thank members of the Wang Laboratory for helpful discussions and technical support. We thank Dr. X. Jia and T.M. Kay for DNA template preparations. This work is supported by the National Institutes of Health grants R01GM136894 (to M.D.W.). M.D.W. is a Howard Hughes Medical Institute investigator. This work was performed in part at the Cornell NanoScale Facility, a member of the National Nanotechnology Coordinated Infrastructure (NNCI), which is supported by the National Science Foundation (Grant NNCI-2025233).

## SUPPORTING MATERIAL

Supporting Methods and four Supplementary Figures are available.

