## Supplementary Information for "Optical Torque Calculations and Measurements for DNA Torsional Studies"

### Numerically calculate the excitation field for FEM simulation

The excitation field for our FEM simulation was calculated via Mathematica and then imported into COMSOL for further simulation. This field calculation was based on the approach described in (1) for a Gaussian (0, 0) mode polarized in x-axis, which can be written as

$$\vec{\boldsymbol{E}}_{\mathrm{ex}}(\rho,\varphi,z) = -\frac{ikfe^{-ikf}}{2}\sqrt{\frac{n_0}{n_1}}\begin{bmatrix} I_{00}+I_{02}\cos(2\varphi) \\ I_{02}\sin(2\varphi) \\ -2iI_{01}\cos\varphi \end{bmatrix}, \tag{S1}$$

$$I_{00} = \int_0^{\theta_{\max}} f_{\mathrm{w}}(\theta)\,(\cos\theta)^{\frac{1}{2}}\sin\theta\,(1+\cos\theta)J_0(k\rho\sin\theta)e^{ikz\cos\theta}d\theta, \tag{S2}$$

$$I_{01} = \int_0^{\theta_{\max}} f_{\mathrm{w}}(\theta)\,(\cos\theta)^{\frac{1}{2}}\sin^2\theta\,J_1(k\rho\sin\theta)e^{ikz\cos\theta}d\theta, \tag{S3}$$

$$I_{02} = \int_0^{\theta_{\max}} f_{\mathrm{w}}(\theta)\,(\cos\theta)^{\frac{1}{2}}\sin\theta\,(1-\cos\theta)J_2(k\rho\sin\theta)e^{ikz\cos\theta}d\theta, \tag{S4}$$

where the coordinate transformation is based on $x=\rho\cos\varphi$ and $y=\rho\sin\varphi$. $k$ is the wave number, $f$ is the focal length, $\theta_{\max}$ is determined by the NA of the objective, $f_{\mathrm{w}}(\theta) = e^{-\frac{1}{f_0^2}\frac{\sin^2\theta}{\sin^2\theta_{\max}}}$, $f_0$ is the aperture filling factor, $n_0$ is the refractive index of air, $n_1$ is the refractive index of glass, and $J_n$ is the $n$th-order of Bessel function.

**Force and torque simulation of a quartz cylinder**

Once the excitation field was imported into COMSOL, it was used to compute the overall E-field distribution within the simulation sphere. For the force simulation on a quartz cylinder, we moved the cylinder axially with its extraordinary axis aligned to the beam polarization, where the permittivity tensor can be described as

$$\overleftrightarrow{\epsilon} = \begin{bmatrix} \epsilon_\mathrm{e} & 0 & 0 \\ 0 & \epsilon_\mathrm{o} & 0 \\ 0 & 0 & \epsilon_\mathrm{o} \end{bmatrix}, \tag{S5}$$

For torque simulation, we rotated the cylinder around its cylindrical axis by angle $\alpha$, where the permittivity tensor became

$$\overleftrightarrow{\epsilon}_\alpha = \begin{bmatrix} \epsilon_\mathrm{e}\cos^2\alpha + \epsilon_\mathrm{o}\sin^2\alpha & (\epsilon_\mathrm{e} - \epsilon_\mathrm{o}) \sin\alpha\cos\alpha & 0 \\ (\epsilon_\mathrm{e} - \epsilon_\mathrm{o}) \sin\alpha\cos\alpha & \epsilon_\mathrm{e}\sin^2\alpha + \epsilon_\mathrm{o}\cos^2\alpha & 0 \\ 0 & 0 & \epsilon_\mathrm{o} \end{bmatrix}. \tag{S6}$$

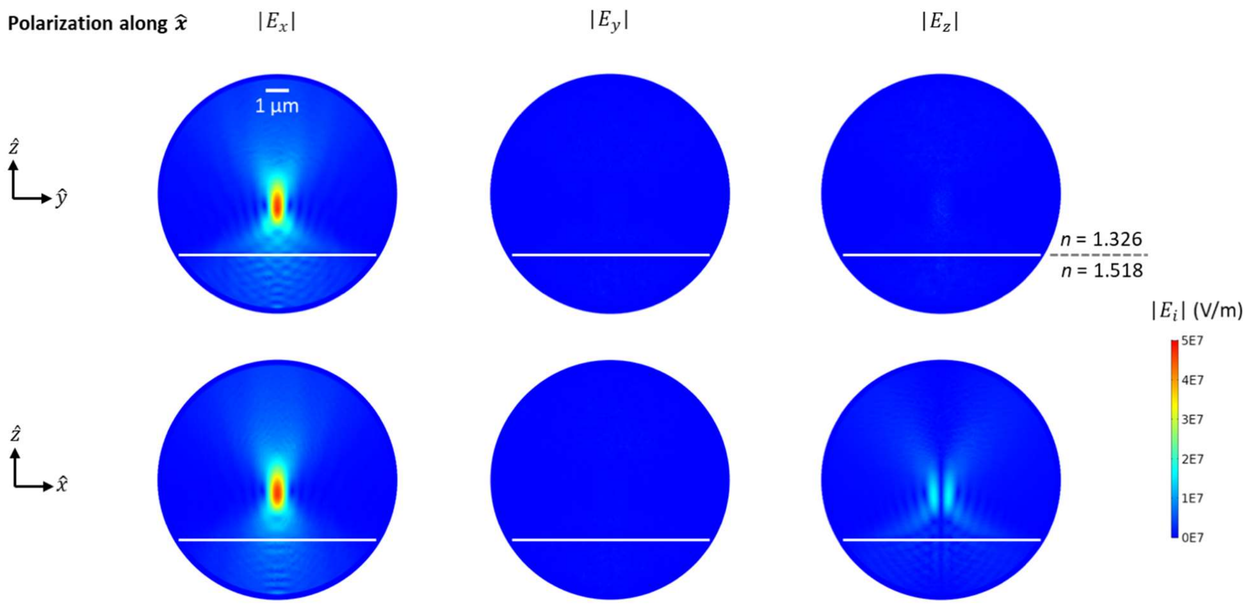


Figure S1. Individual components of simulated E-field distribution of the focused beam with in index mismatch as shown in Fig. 2c.

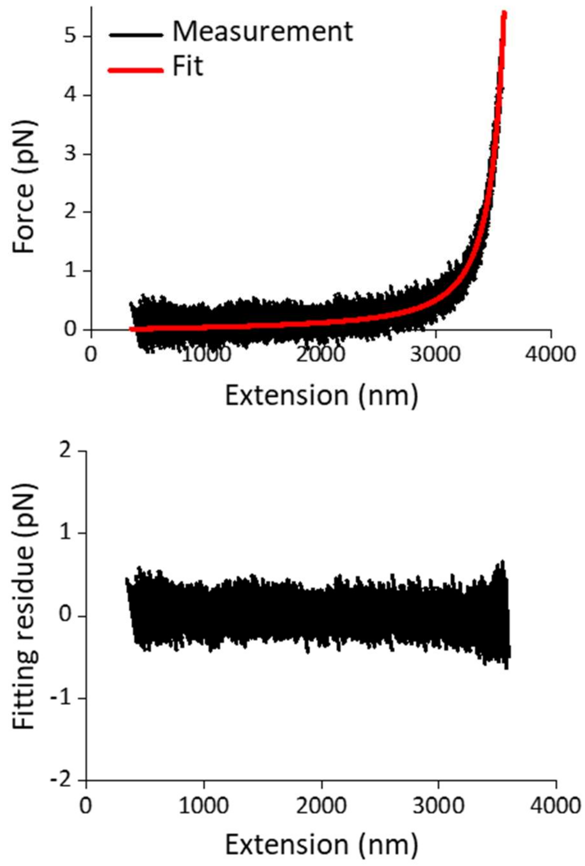


Figure S2. DNA elastic property measurements. The top panel shows an example trace of the 11.5 kb DNA force-extension measurement of a single trace and its fit by the Marko-Siggia worm-like chain model (2). The bottom panel shows the residue of the fit. The fitting to traces such as the one shown here yields a DNA persistence length of 46.0 $\pm$ 2.6 nm (mean $\pm$ SD, $N$ = 13 traces), consistent with previously published results (3). In addition the number of base pairs from the fit is 11,361 ± 112 bp (mean $\pm$ SD), while the expected number of base pairs is 11,516 bp.

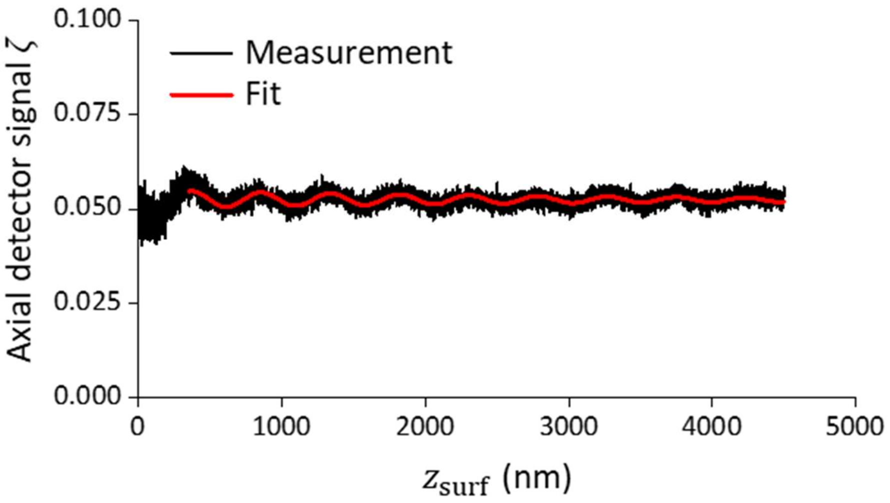


Figure S3. An example trace of the focal shift ratio $f_{\mathrm{s}}$ determination based on the Fabry-Pérot effect. The signal was fit by $\zeta = a_1 + a_2 \cdot z_{\mathrm{surf}} + a_3 \sin\left(\frac{4\pi n_{\mathrm{w}}}{\lambda_0} \cdot f_{\mathrm{s}} z_{\mathrm{surf}} + a_4\right) \cdot e^{-a_5 \cdot z_{\mathrm{surf}}}$, where $\lambda_0$ = 1064 nm is the wavelength of the light in vacuum and $n_{\mathrm{w}}$ is the refractive index of aqueous solution. The fitting to the period of the interference pattern gives $f_{\mathrm{s}}$ (4). Here, we measured *N* = 16 cylinders, resulting in $f_{\mathrm{s}}$ = 0.829 $\pm$ 0.003 (mean $\pm$ SD). This value is ~ 2% smaller than that obtained using the dsDNA ruler method (Fig. 4b).

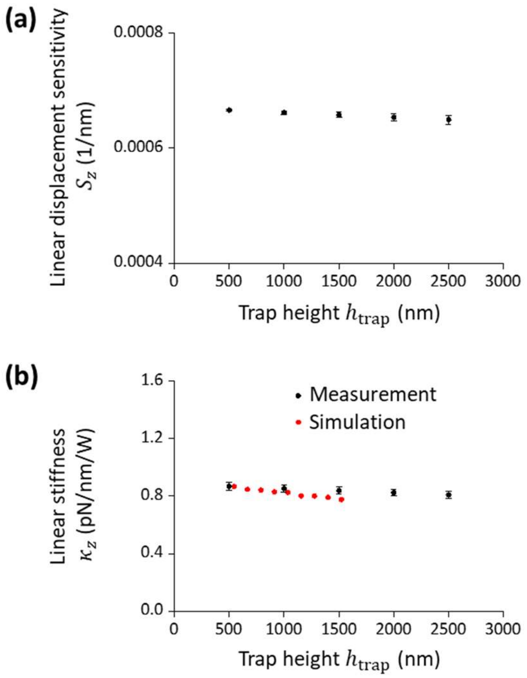


Figure S4. Experimental determination of the axial position detector sensitivity and axial linear trap stiffness as a function of the trap height, using the DNA unzipping method (5).

(a) Axial position detector sensitivity as a function of the trap height. Error bars are SDs from $N$ = 6 cylinders.

(b) Axial linear stiffness as a function of the trap height. Error bars are SDs from $N$ = 6 cylinders. For comparison, the results from the simulations are also shown.
